# Activation of the *Pseudomonas aeruginosa* Glycerol Regulon Promotes Antibiotic Persistence and Modulates Virulence Phenotypes

**DOI:** 10.1101/2025.04.14.648800

**Authors:** Nicholas Evans, Jessica A. Scoffield

**Affiliations:** Department of Microbiology, University of Alabama at Birmingham, Birmingham AL, United States

## Abstract

Chronic infections with *Pseudomonas aeruginosa* are a major contributor of lung decline in persons with cystic fibrosis (pwCF). *P. aeruginosa* establishes life-long infections in the CF airway by utilizing various adaptation strategies to persist, including altering the expression of metabolic genes to acquire nutrients that are abundant in the CF airway. Glycerol, which is readily available in the airway, is imported and metabolized by genes in the *glp* regulon, which is under the control of the GlpR repressor. Previously, it has been shown that the loss of GlpR results in increased biofilm development in a CF-adapted isolate of *P. aeruginosa* compared to a wound isolate. Based on the increased biofilm phenotype previously observed and because biofilms are associated with increased antibiotic tolerance, we questioned whether GlpR plays a role in mediating antibiotic resistance of *P. aeruginosa*. In this report, we show that loss of GlpR increases tobramycin resistance of a CF-adapted isolate in synthetic sputum and in airway epithelial cell and *Drosophila melanogaster* colonization models. Further, transcriptomics analysis revealed that CF-adapted mutants of *glpR* overexpresses genes involved in multidrug tolerance and chronic infection phenotypes such as alginate. In summary, our study illustrates that activation of the glycerol (*glp*) regulon may promote *P. aeruginosa* persistence in the CF airway.

## INTRODUCTION

Chronic *Pseudomonas aeruginosa* airway infections are the leading cause of morbidity and lung decline in persons with cystic fibrosis (pwCF) (1). CF is the most common lethal genetic disorder in individuals of European descent and is caused by mutations in the cystic fibrosis transmembrane conductance regulator channel (CFTR) protein. As an anion channel, CFTR maintains ion balance within the airway, which facilitates clearance of mucus and promotes airway hydration (2). However, loss of CFTR leads to impaired immune function, reduced mucociliary clearance, and the accumulation of thick mucus in the lungs, which support the chronic colonization of diverse microbes (3).

*Pseudomonas aeruginosa* is the dominant airway pathogen in adults with CF and is associated with worsening lung function and declining microbial diversity (4, 5). The success of *P. aeruginosa* as a chronic pathogen in the airway is primarily due to its intrinsic drug resistance, ability to modify the regulation of virulence and metabolic genes, conversion to the mucoid phenotype, and ability to form recalcitrant biofilms to evade the host immune response and antimicrobials (1, 4–6). Although the introduction of highly effective modulator therapy (HEMT) to correct CFTR function has remarkably improved the quality of life of pwCF and transformed the landscape of CF intervention, *P. aeruginosa* persists after HEMT (7–9), suggesting that post-HEMT conditions in the CF airway still support a chronic lifestyle of growth for *P. aeruginosa*.

Host-derived nutrients are mediators of microbial colonization, pathogenesis, and microevolution of *P. aeruginosa.* Airway sputum supports the growth and nutritional requirements of *P. aeruginosa* during CF infection and modulates the expression of genes that contribute to persistence and virulence determinants that are critical for chronic infection (6, 10–12). CF sputum is composed of both host-derived and bacterial-related products, including the major lung surfactant, phosphatidylcholine (PC). Phospholipase C produced by *P. aeruginosa* cleaves PC into phosphorylcholine, fatty acids, and glycerol, and as a result, liberated glycerol can be used as a potential nutritional source for *P. aeruginosa* (13, 14). Previous studies have demonstrated that genes (*glpD* and *glpK*) specific to the *glp* (glycerol) regulon are constitutively expressed in some *P. aeruginosa* CF isolates recovered from sputum and are required for *in vivo* degradation of PC. The *P. aeruginosa glp* (glycerol) regulon encodes a membrane-associated glycerol diffusion facilitator (GlpF), a glycerol kinase (GlpK), a membrane protein involved in alginate biosynthesis (GlpM), and a glycerol-3-phosphate dehydrogenase (GlpD). Host-derived glycerol is transported into the cell via GlpF and is phosphorylated to glycerol 3-phosphate (G3P) by GlpK, whereas exogenous glycerol 3-phosphate is transported into the cell via the GlpT transporter system (15). Importantly, G3P induces the *glp* regulon by binding to GlpR, the *glp* repressor (15). We have previously shown that glycerol metabolism and loss of the *glp* regulon transcriptional repressor, GlpR, promotes biofilm development by *P. aeruginosa* in a CF-adapted isolate, which is caused by the overproduction of Pel exopolysaccharide (16). Pel is a cationic exopolysaccharide structural component of the biofilm matrix that contributes to aminoglycoside resistance (17–19). Biofilm development within the CF airway is correlated with increased persistence and the establishment of a chronic infection that contributes to *P. aeruginosa* tolerance to antimicrobials (20, 21). Hence, we questioned the contribution of the glycerol regulon in *P. aeruginosa* persistence to the CF airway.

In this study, we tested whether GlpR mediates *P. aeruginosa* persistence during exposure to the aminoglycoside tobramycin, an antibiotic commonly used to treat chronic *P. aeruginosa* airway infections (22–25). We report that loss of GlpR promotes tolerance to tobramycin in a CF-adapted strain of *P. aeruginosa* on CF and non-CF airway epithelial cells, and in a *Drosophila melanogaster* model of infection. Further, transcriptomics analysis revealed that GlpR modulates the expression of genes involved in antibiotic tolerance. Finally, we show that loss of GlpR promotes the production of exopolysaccharide alginate production, which is associated with chronic CF airways infections, but decreases the production of pyocyanin, a secreted product that is generally produced at higher levels during an acute infection. Taken together, our data demonstrate that the *P. aeruginosa* response to host-derived glycerol, which is controlled by the GlpR repressor, may facilitate persistence and adaptation to chronic in the CF airway.

## RESULTS

### Loss of GlpR Promotes *P. aeruginosa* Resistance to the Frontline CF therapeutic Tobramycin

*P. aeruginosa* liberates glycerol from host-derived surfactant during colonization in the CF airway (14). Further, genes involved in the glycerol regulon (*glp*) have been shown to be upregulated in *P. aeruginosa* isolates recovered from CF sputum (6), suggesting that glycerol may be an important nutrient within the CF airway. Additionally, our previous studies demonstrate that glycerol metabolism promotes biofilm development in a CF-adapted *P. aeruginosa* isolate. In this report, we tested whether glycerol metabolism via the loss of the *glp* regulon repressor, GlpR, mediates CF-adaptive phenotypes that mediate chronic infection and persistence in the airway. First, we tested the loss of GlpR on antibiotic susceptibility in a non-CF (PAO1) and CF-adapted (FRD1) isolate of *P. aeruginosa* using three classes of antibiotics, including an aminoglycoside (tobramycin), cephalosporin (ceftazidime), quinolone (ciprofloxacin), which inhibit protein synthesis, cell wall synthesis, and DNA replication, respectively. Loss of *glpR* promoted both PAO1 and FRD1 survival in the presence of tobramycin, but not ceftazidime and ciprofloxacin compared to wild-type, as indicated by a higher minimal inhibitory concentration (MIC) required to target the *glpR* mutants (Figure 1A-1C). Next, we tested whether this resistance was conserved during exposure to other aminoglycosides (streptomycin, kanamycin, and amikacin) and observed that loss of *glpR* increased resistance of FRD1 to amikacin, but not PAO1 (Figure S1). Further, a defect in *glpR* did not impact FRD1 or PAO1 during exposure to streptomycin or kanamycin (Figure S1). Tobramycin and amikacin are two preferred aminoglycosides used to target chronic *P. aeruginosa* infections in the CF airway (22, 26). Taken together, our data illustrate that glycerol metabolism potentially facilitates the persistence of *P. aeruginosa* during antibiotic therapy.

**Figure 1:**
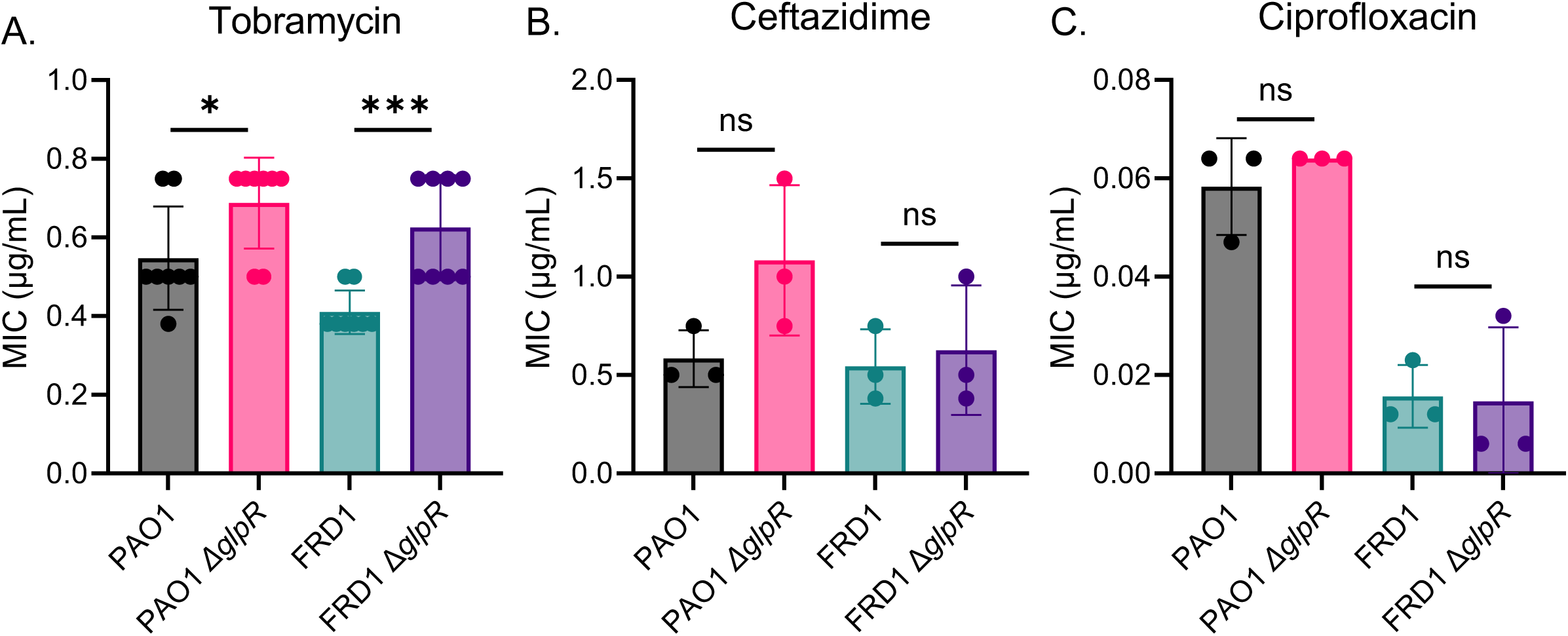
Loss of *glpR* promotes *P. aeruginosa* tolerance to tobramycin. Determination of MIC for three drug classes with *glpR* knockout. All MIC assays were conducted using Mueller Hinton Agar with lawns grown 24 hours, and MIC strips provided by Liofilchem. MIC measurements for strains exposed to a gradient of A). the aminoglycoside tobramycin (n=8) B). the beta lactamase ceftazidime (n=3) C). the fluoroquinolone ciprofloxacin (n=3) *Statistical significance assessed by t-test. *P<0.05, ***P<0.001

### The CF-adapted isolate, FRD1, displays enhanced antibiotic resistance to tobramycin in CF epithelial cell and *Drosophila melanogaster* colonization models

Our initial observations indicated that *P. aeruginosa glpR* mutant strains confer resistance to tobramycin, a frequently used antibiotic therapy for *P. aeruginosa* CF infection. To further examine the tobramycin resistance phenotype in a CF-relevant condition, we grew PAO1, FRD1, and the *glpR* mutants in SCFM2, a synthetic sputum media that closely recapitulates the nutritional environment of the CF airway. As shown in Figure 2A, the PAO1 Δ*glpR* mutant displayed enhanced exponential and biphasic growth during enhanced in the presence of 5 ug/mL of tobramycin compared to wildtype PAO1, but demonstrated comparable growth at the end of 36 hours. however, both the wildtype and Δ*glpR* mutant strain were both inhibited during growth on 10 ug/mL of tobramycin. In contrast, the FRD1 Δ*glpR* mutant displayed a significant growth advantage in the presence of 10 ug/mL of tobramycin compared to wildtype FRD1 (Figure 2B). There were no significant differences between FRD1 and the FRD1 Δ*glpR* mutant at 5 ug/mL of tobramycin. It is also important to note that there are no growth differences between PAO1, FRD1, and their respective *glpR* mutants without tobramycin treatment (Figures 2A and 2B).

**Figure 2:**
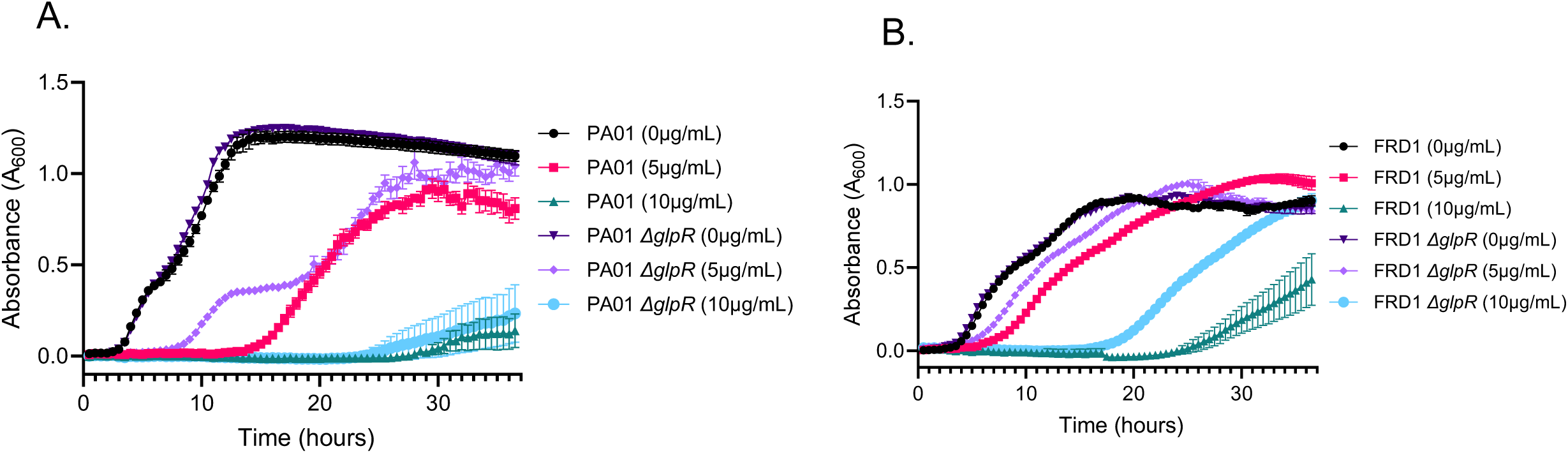
Loss of *glpR* promotes tobramycin resistance in the CF-adapted isolate in SCFM2. A). Growth curves of PAO1 and PAO1 *ΔglpR* grown in synthetic cystic fibrosis sputum media 2, with or without 5 and 10 μg/mL tobramycin. (n=3); B) Growth curves of FRD1 and FRD1 *ΔglpR* grown in SCFM2, with or without 5 and 10 μg/mL tobramycin. (n=3)

Although SCFM2 mimics the CF lung nutritional environment (27), we also wanted to validate our tobramycin tolerance findings using an epithelial cell infection model and *Drosophila melanogaster in vivo* colonization model. Initial colonization of PAO1, FRD1, and the *glpR* mutants showed no differences in the initial colonization of wildtype (16HBEs) and CF (CFBEs) epithelial cells by wildtype *P. aeruginosa* or the *glpR* mutants. Further, there were no differences in cytotoxicity in the wildtype and *glpR* mutant strains as indicated by the LDH assay (Figures 3A and 3B) prior to tobramycin treatment. Consistent with our previous findings, the FRD1 Δ*glpR* mutant demonstrated increased tolerance to 20 µg/mL of tobramycin compared to wildtype FRD1 in both 16HBEs and CFBEs. In contrast, the PAO1 *glpR* mutant did not promote tolerance to tobramycin compared to wildtype PAO1. Moreover, cells infected with wildtype and *glpR* mutant *P. aeruginosa* and treated with tobramycin did not exhibit any differences in cytotoxicity (Figures 3C and 3D). Lastly, FRD1 and FRD1 Δ*glpR* mutant colonization in the *D. melanogaster* colonization model also showed that loss of *glpR* increases resistance to tobramycin (Figure 4). In sum, these data indicated that activation of the *glp* regulon by host-derived glycerol may facilitate antibiotic persistence in the CF airway.

**Figure 3:**
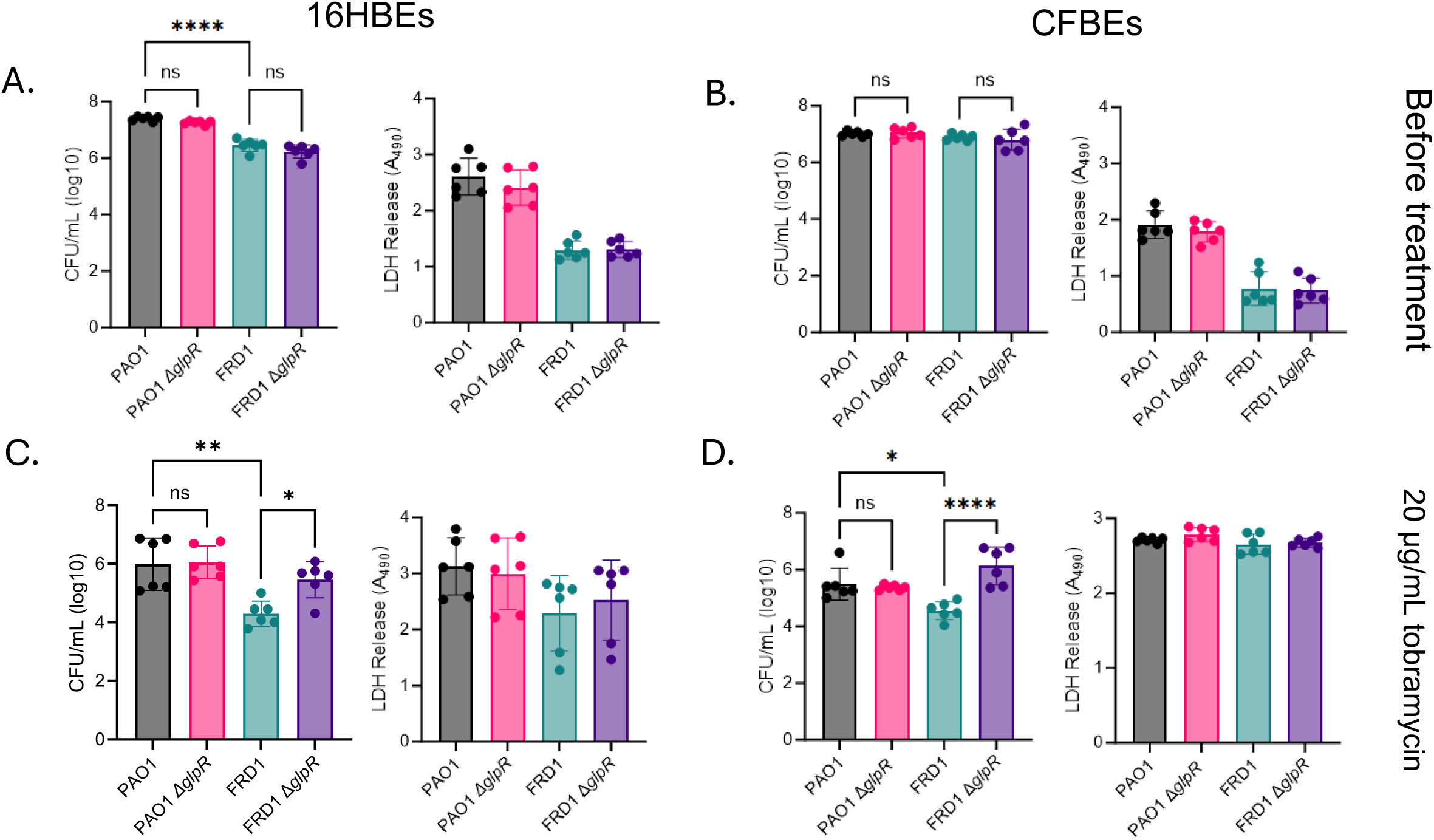
A defect in *glpR* enhances tobramycin persistence in an epithelial cell model in the CF-adapted isolate FRD1. Colony forming units and cytotoxicity of PAO1, FRD1, and the *glpR* mutants in 16HBEs (A) and CFBEs (B) prior to tobramycin treatment. Colony forming units and cytotoxicity of PAO1, FRD1, and the *glpR* mutants in 16HBEs (C) and CFBEs (D) post tobramycin treatment. (n=3) *Significance by one-way ANOVA. *P<0.05, **P<0.01, ****P<0.0001

**Figure 4:**
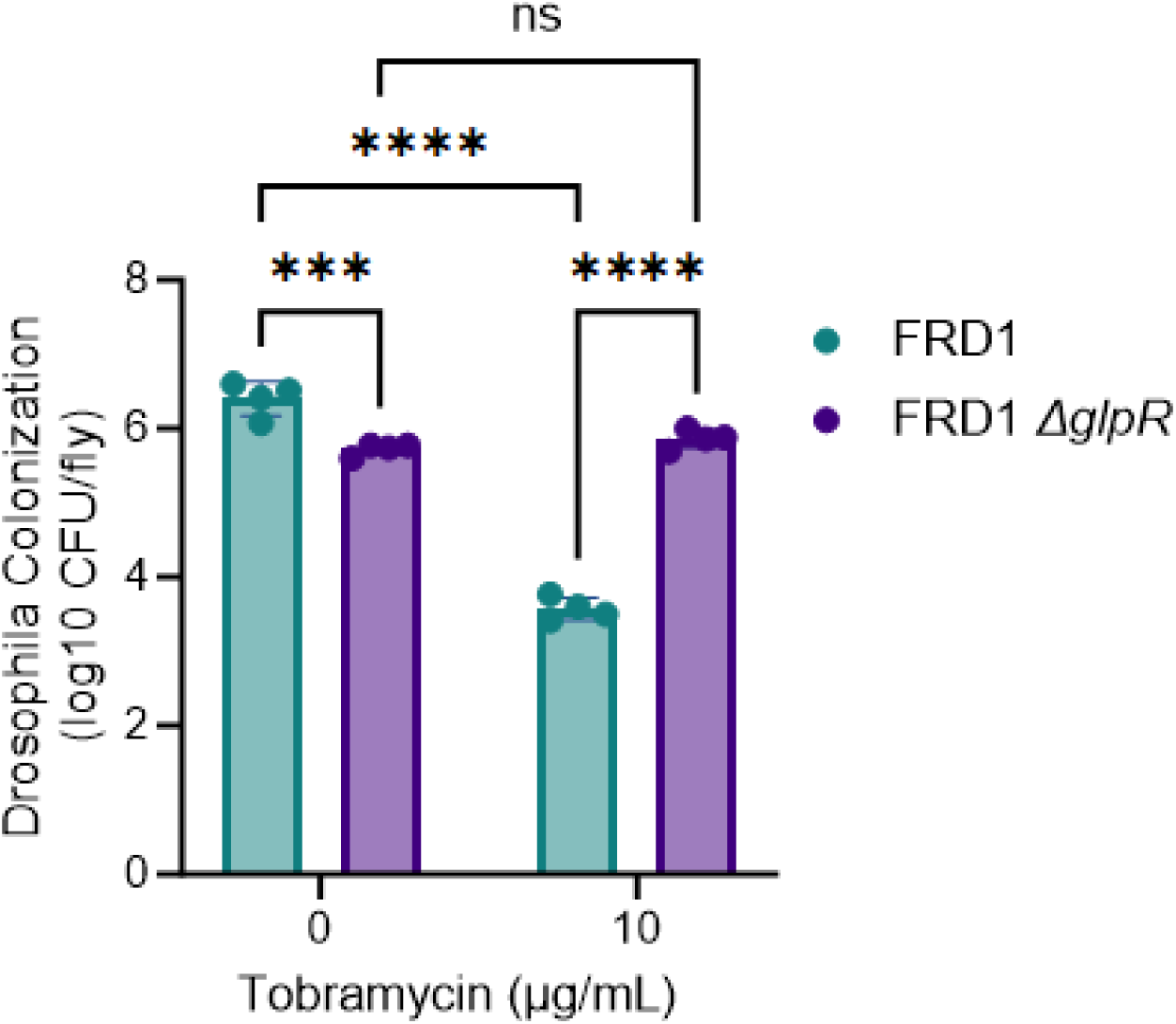
Loss of *glpR* promotes tobramycin tolerance in *Drosophila melanogaster* in the CF-adapted isolate FRD1. Data represent 4 biological replicates, with 10 flies per group for each replicate. *Statistical significance determined by two-way ANOVA on log transformed data. ns= not significant, ***P<0.001, ****P<0.0001.

### Transcriptomics Reveals that GlpR Modulates the Expression Antibiotic Tolerance Genes

In an effort to determine molecular mechanisms that contribute to how the *glp* regulon mediates increased tolerance to antimicrobials in FRD1, we examined the transcriptomes of wildtype and *glpR* mutant PAO1 and FRD1 cultures, with and without tobramycin treatment, to identify overexpressed antibiotic responsive genes solely in the FRD1 Δ*glpR* background. As expected, we observed increases in gene expression in the Mex multidrug efflux system with tobramycin treatment in both the PAO1 and FRD1 wildtype backgrounds, as well as in the PAO1 Δ*glpR* and FRD1 Δ*glpR* mutants in response to treatment (Figure 5A and 5B). When we compared PAO1 to PAO1 Δ*glpR* with treatment, antibiotic tolerance genes were downregulated or unchanged in the *glpR* mutant. Notably, when we compared FRD1 and FRD1 Δ*glpR* with tobramycin treatment, most genes were either downregulated or unchanged in the *glpR* mutant, with the exception of *mexA*, *carO,* and PA5159, which were all upregulated (Figure 5B). MexA is a component of the MexAB-OprM efflux pump apparatus that contributes to drug resistance in *P. aeruginosa* (28). CarO is a periplasmic protein that is predicted to be involved in calcium homeostasis and antibiotic tolerance (29). PA5159 is a multidrug resistance protein located in the cytoplasmic membrane that is in an operon with PA5160, a drug efflux protein (30). The overexpression of these genes strictly in the FRD1 Δ*glpR* mutant background compared to wildtype FRD1, PAO1, or the PAO1 Δ*glpR* mutant in the presence of tobramycin indicates that activation of the *glp* regulon may uniquely modulate tolerance in the CF-adapted isolate FRD1, but not PAO1.

**Figure 5:**
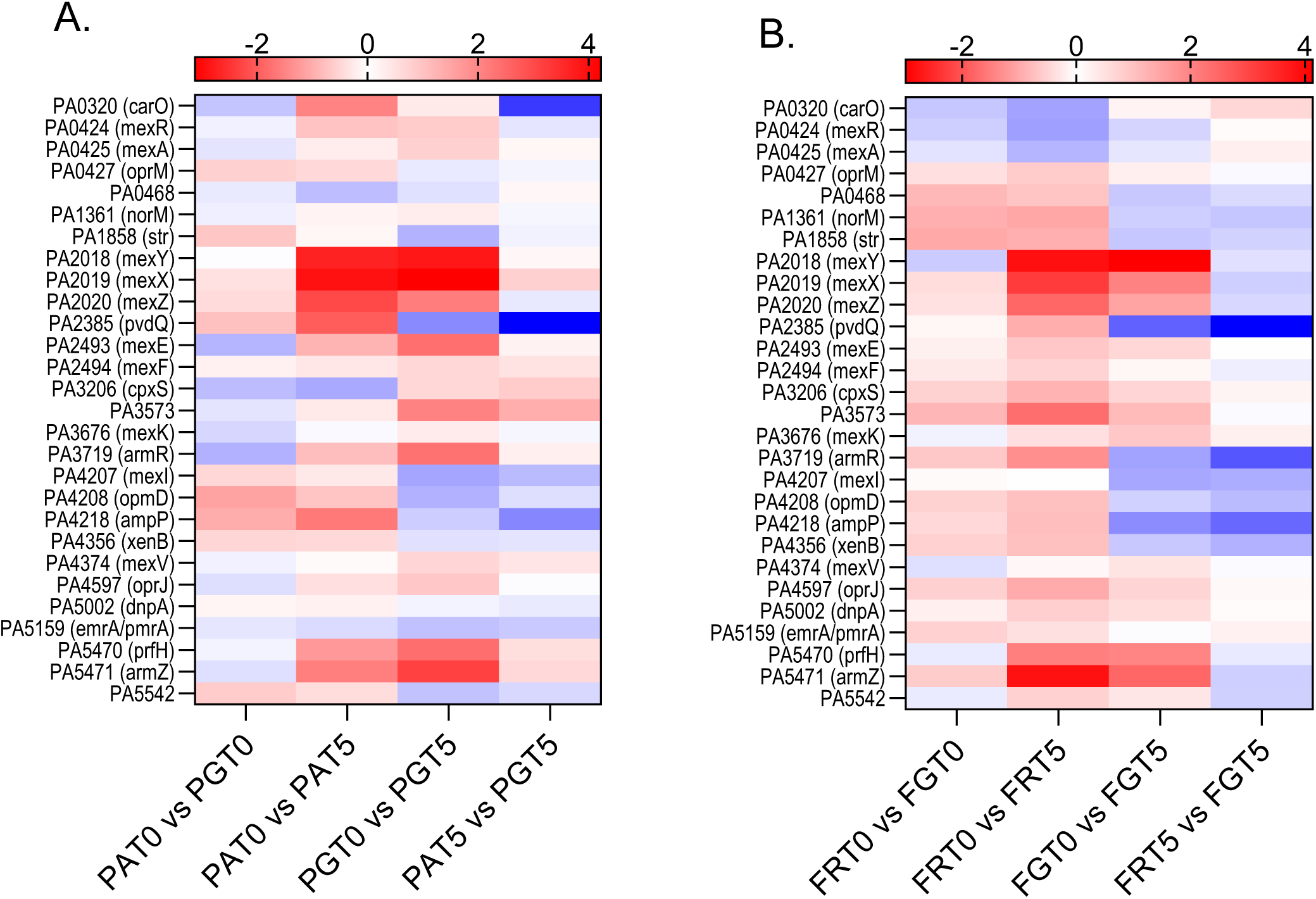
Transcriptomic changes in response to tobramycin and loss of *glpR*. RNA-Seq results were narrowed by genes in the KEGG pathway for response to antibiotic (GO:0046677), then further trimmed for genes where at least one result had a significant p-value. Log fold changes are color coordinated from greatest (red) to least (blue). A) Comparisons between PAO1 (PA) and PAO1 *ΔglpR* (PG) with and without treatment of 5 μg/mL tobramycin (T5) in SCFM2. B) Comparisons between FRD1 (FR) and FRD1 *ΔglpR* (FG) with and without treatment of 5 μg/mL tobramycin (T5) in SCFM2. RNA sequencing was performed in triplicate.

### Loss of GlpR mediates the production of virulence determinants involved in chronic infection

We observed phenotypic differences between the wildtype and mutant strains. Hence, we measured the production of a subset of virulence factors in the wildtype and *glpR* mutants that are significant to CF airway disease. Pyocyanin, a blue-green phenazine pigment that contributes to cell toxicity (31), was reduced in the FRD1 Δ*glpR* strain compared to wildtype and restored in the *glpR* complemented strain (Figure 6A). In contrast, pyocyanin production was increased in the PAO1 Δ*glpR* strain compared to wildtype and restored to basal levels in the complemented strain (Figure 6B). In support of these findings, transcriptomics analysis showed the overexpression of three phenazine biosynthetic genes, *phzC1*, *phzE1*, and *phzG1* in the PAO1 Δ*glpR* background compared to wildtype PAO1, and the downregulation of *phzB2* in the FRD1 Δ*glpR* background (Table 1).

**Figure 6:**
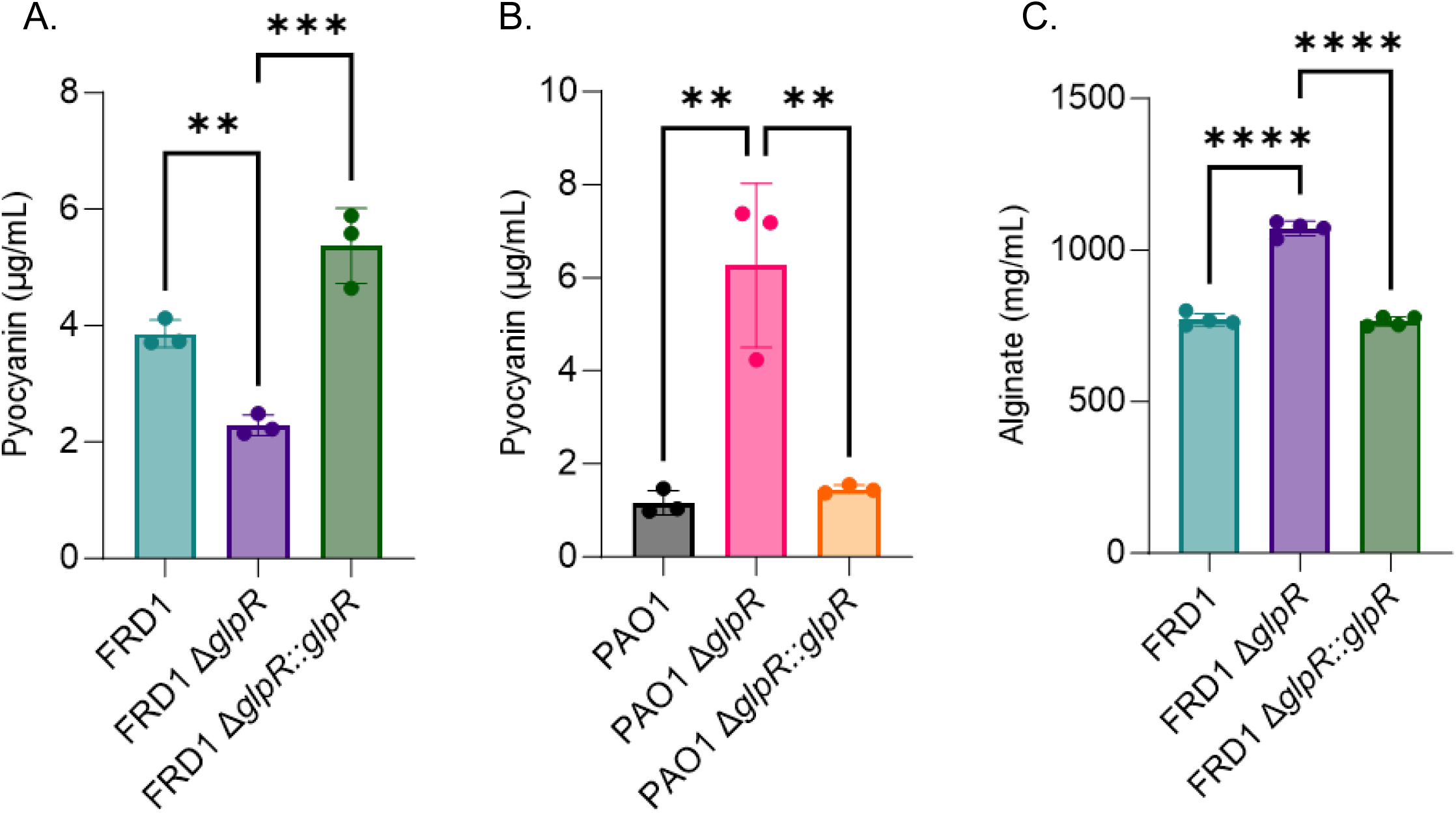
GlpR modulates the production of virulence factors. Cultures were grown in SCFM2 for 24 hours prior to measuring pyocyanin and alginate. A. Pyocyanin production in FRD1, FRD1 Δ*glpR*, and FRD1 Δ*glpR*::*glpR* complemented strains. B. Pyocyanin production in PAO1, PAO1 Δ*glpR*, and PAO1 Δ*glpR*::*glpR* complemented strains. C. Alginate production in FRD1, FRD1 Δ*glpR*, and FRD1 Δ*glpR*::*glpR* complemented strains. *Significance by one-way ANOVA. **P<0.01, ***P<0.001, ****P<0.0001

**Table 1:**
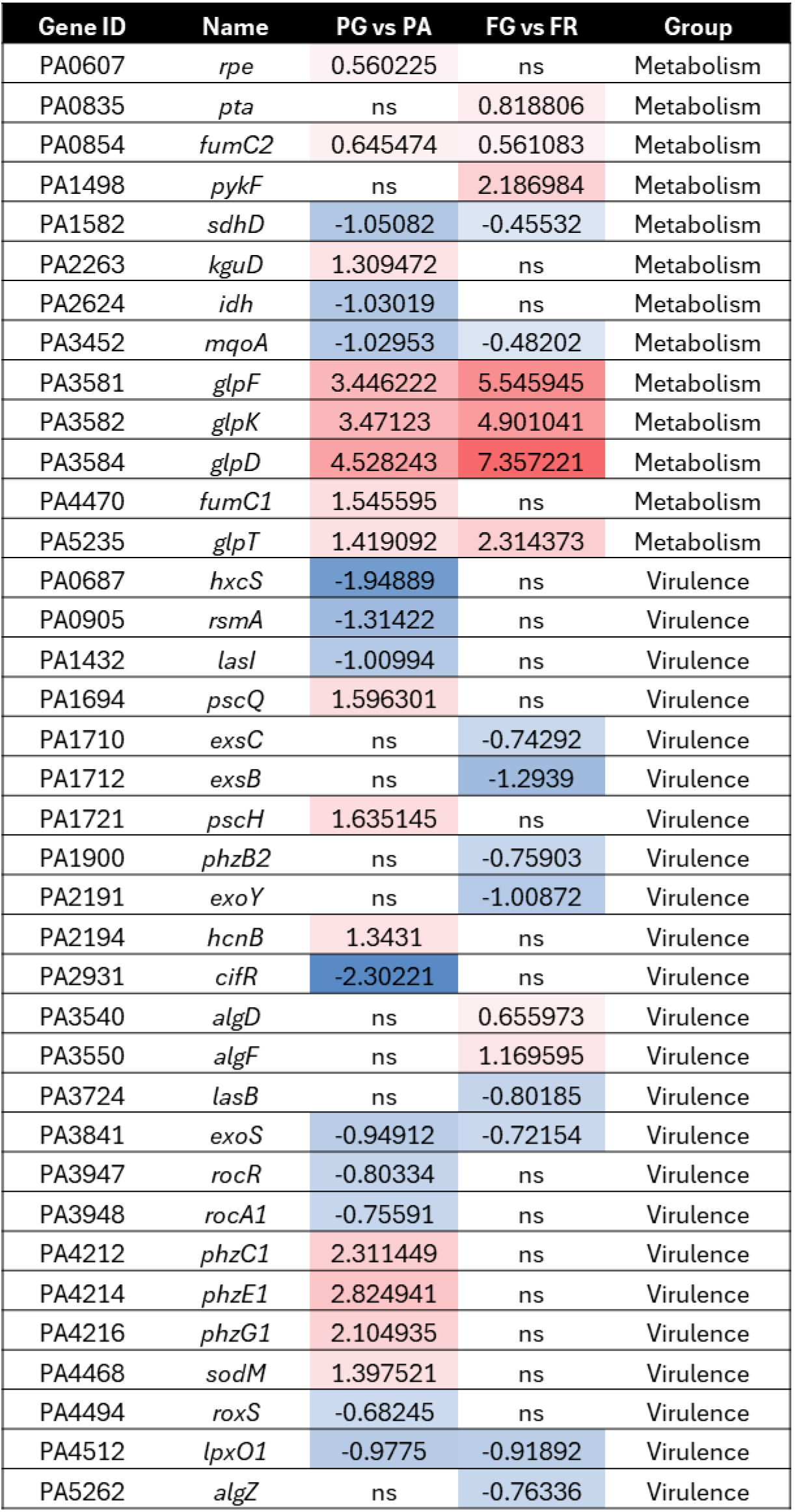
Differential expression of metabolic and virulence related genes in the wildtype vs *glpR* mutants. Log fold changes of genes with significant differences in expression are shown in column for PAO1 vs PAO1 *ΔglpR* and FRD1 vs FRD1 *ΔglpR.* All strains were grown in SCFM2 for RNA sequencing and are the results of three biological replicates (n=3). ns = not significant

The hallmark of a chronic *P. aeruginosa* infection in the CF airway is the conversion to the mucoid phenotype, which is caused by an overproduction of alginate (32). The CF-adapted isolate, FRD1, displays mucoidy while the wound isolate, PAO1, does not, therefore, we measure alginate production in FRD1 and the FRD1 Δ*glpR* mutant. As shown in Figure 6C, loss of *glpR* promoted alginate production compared to wildtype FRD1, and this increase was abolished in the complemented strain. In agreement with these results, the alginate biosynthetic genes *algD* and *algF,* were overexpressed in the FRD1 Δ*glpR* background in the transcriptomics analysis (Table 1).

## DISCUSSION

Nutrient availability and acquisition are key drivers of *P. aeruginosa* adaptation and evolution in the CF airway (10–12, 33). The ability to metabolize host-derived nutrients signals specific bacterial responses that influence the differential expression of virulence genes or pathogenicity (33, 34). In the CF lung, glycerol is a major host-derived nutritional source for *P. aeruginosa*, as evident by the constitutive expression of *glp* genes in *P. aeruginosa* recovered from CF sputa (6). In previous studies, we observed that glycerol metabolism promotes biofilm development by *P. aeruginosa* (16), which is the main mode of bacterial growth in the CF airway during chronic infection (19, 22, 32, 35–37).

Specifically, we found that the increase in biofilm development by the CF-adapted isolate in a *glpR* deficient CF-adapted isolate was due to the overproduction of the exopolysaccharide Pel (16), which has been shown to mediate tolerance to antimicrobials (17, 19, 21, 36, 38). Due to these findings, we questioned whether activation of the glycerol metabolic genes modulates antibiotic persistence and additional chronic infection phenotypes. In this study, we examined the role of the *glp* regulon repressor, GlpR, on antibiotic persistence, colonization in an epithelial cell and *Drosophila* model, and virulence production. Our findings illustrated that loss of *glpR* promoted tolerance to the frontline CF antibiotic tobramycin in the CF-adapted isolate FRD1 in SCFM2, wildtype and CF epithelial cells, and in a Drosophila colonization model. Transcriptomics analysis revealed the overexpression of three antibiotic resistance genes (*mexA*, *carO*, and PA5159) in the FRD1 *glpR* defective strain. Moreover, loss of *glpR* facilitated the overproduction of alginate, but reduction of pyocyanin in the FRD1 background, but pyocyanin was increased the PAO1 *glpR* mutant. Overproduction of alginate and downregulation of toxins, such as pyocyanin, are hallmarks of chronic infection in the CF airway, although pyocyanin production can be variable among CF isolates. Overall, these findings indicate that glycerol metabolism may regulate the switch from an acute to chronic infection lifestyle in *P. aeruginosa*.

Glycerol metabolism has been described in other bacteria to be a critical mediator of persistence during infection. For example, *Borrelia burgdorferi* utilizes host-derived glycerol and the *glp* operon to persist in ticks (39, 40). Additionally, loss of glycerol-3-phosphate dehydrogenase (*glpD*) reduces persister cell formation in the marine pathogen *Vibrio splendidus* (41). Strikingly, glycerol metabolism mediates intracellular survival and persistence by *Listeria monocytogenes* and *Mycoplasma pneumoniae* (42–46). Similarly, glycerol metabolism contributed to persistence of a CF-adapted isolate in our study. We observed that loss of GlpR increased FRD1’s resistance to tobramycin in an artificial sputum media (SCFM2), on airway epithelial cells, and in a Drosophila colonization model, whereas the PAO1 *glpR* mutant strain was only slightly able to persist at a lower concentration of tobramycin in SCFM2, but not on epithelial cells. Previous work by our group and others has shown that the expression of *glp* genes and glycerol metabolism promotes the overproduction of the exopolysaccharide Pel (16, 47). We have previously shown that loss of GlpR results in the upregulation of *pelA* expression and overproduction of Pel in the CF-adapted isolate FRD1, but not in the wound isolate, PAO1. Pel is a cationic polysaccharide that is important for cell-to-cell interactions within *P. aeruginosa* biofilms and promotes tolerance to the aminoglycoside tobramycin (17, 18, 48–52). Based on our previous and current studies, in addition to work performed by other groups, we speculate that increased tolerance to tobramycin by FRD1 in SCFM2, in addition to increased persistence in response to tobramycin in the epithelial cell and Drosophila colonization models is partially due to the overproduction of Pel. Additionally, we observed that loss of *glpR* promoted alginate production and expression of three antibiotic resistance genes, *carO*, *mexA*, and PA5159. Similar to Pel, the overproduction of alginate is associated with increased tolerance to antimicrobials and the host immune response (19, 21, 32, 38). MexA is a component of the MexAB-OprM multidrug efflux pump system (28). In *Acinetobacter baumannii*, CarO, an outer membrane protein, has been shown to alter membrane permeability to facilitate resistance to carbapenem (29), however the role of *carO* in drug tolerance in *P. aeruginosa* is now well understood. Similarly, although not well studied, PA5159, encodes a predicted multidrug transporter (30). Taken together, our findings illustrate that activation of the *glp* regulon may promote drug tolerance in CF-adapted *P. aeruginosa* by enhancing the production exopolysaccharides like alginate and increasing the expression of antibiotic tolerance genes.

The switch from an acute to chronic lifestyle of infection by *P. aeruginosa* in the CF airway is characterized by biofilm development, drug resistance, alginate production, and a decrease in toxin production (5) Collectively, our studies on glycerol metabolism indicate that GlpR enhances biofilm development, antibiotic tolerance, Pel, and alginate production, but decreases the production of the toxin pyocyanin, although pyocyanin production is variable among CF-adapted isolates of *P. aeruginosa.* These findings suggest that the liberation and acquisition of host-derived glycerol by *P. aeruginosa* may facilitate persistence and chronic infection in the CF airway.

In summary, our study shows that a defect in GlpR, the repressor of the glycerol regulon, stimulates chronic infection phenotypes, including antibiotic tolerance and alginate production in a CF-adapted isolate of *P. aeruginosa*. The microevolution exhibited by *P. aeruginosa* CF isolates has proven to be a successful strategy not only to evade the host immune response and avoid clearance from the lung but also to utilize readily available nutrients present in the host environment and persist. Our findings indicate that the acquisition of host-derived glycerol may act as a metabolic signal for persistence. These findings are particularly important because recent studies have shown that CFTR modulator therapy does not completely eradicate *P. aeruginosa* from the lung (7–9, 53). Thus, targeting glycerol metabolic genes may be beneficial for enhancing the clearance of *P. aeruginosa*. Future studies will dissect the regulatory roles of GlpR on antibiotic resistance and genes that contribute to biofilm development.

## MATERIALS AND METHODS

### Bacterial strains and culture conditions

The wound-derived lab strain PA01 and CF-adapted lab strain FRD1 were used as parent strains in this study in addition to isogenic *glpR* mutants in each background and the *glpR* complemented mutants (16). Overnight cultures were grown in lysogeny broth (LB) in a 37°C shaker. Overnight cultures were subcultured in fresh LB until A_600_ reached ∼0.6-0.8 for all experiments.

### *In vitro* Antibiotic Testing

To assess sensitivity of strains to antibiotics, cultures were diluted as described above with the addition of 5 or 10 µg/mL of tobramycin and grown for 30 hours in a BioTek Synergy HTX multi-mode reader. Absorbance (*A*_600_) was measured after 24 hours or over 36 hours with 30-minute intervals for growth curves. Cultures were grown in microtiter plates containing lysogeny broth, Synthetic Cystic Fibrosis Medium (SCFM2) (SynthBiome), or Mueller Hinton broth. Additionally, Mueller Hinton Agar was used to determine the minimal inhibitory concentration of tobramycin using antibiotic gradient strips. Plates were incubated at 37°C for 24 hours before results were read.

### Air-Liquid Interface Cell Culture

Two human bronchial epithelium cell lines were used for this work, the 16HBE cell line was used as a wildtype, or healthy control and cystic fibrosis bronchial epithelial cells, CFBE4lo- (CFBE) (22). Cell lines were maintained in T75 flasks with Minimal Essential Media (MEM) supplemented with 10% FBS (Fetal Bovine Serum) and 100 IU/mL penicillin-streptomycin and grown at 37°C with 5% CO_2_. For air liquid interface culture, 12-well transwell plates were seeded with 5×10^6^ 16HBE or CFBE cells per well and grown submerged in MEM with 10% FBS until confluent. The apical media was then removed, and cells were allowed to polarize at least 7 days before use. For infections, overnight cultures were diluted to A_600_=0.5, which yielded 1×10^6^ CFU/mL. One hour prior to infection, the media in the transwells were replaced with MEM without FBS and the cells are given an hour to adjust. *P. aeruginosa* culture (5µL) was added to the apical media, the plate was shaken gently, and the infected epithelia cells were transferred to a non-sterile incubator to attach and grow for one hour. After the initial hour, the apical media was replaced with 500 μL of fresh MEM and the plate was returned to the incubator for 5 more hours. For plates receiving tobramycin treatment, the media in the transwells was removed and replaced with MEM + 20 ug/mL tobramycin. The plates were then placed back in the incubator for 21 hours based on the established protocol used by the O’Toole lab. To enumerate bacteria colony forming units (CFUs), 500 μL of PBS was added apically and the cells were scraped from the transwell using 1mL pipette tips, then collected in a 1.5 mL conical tube. Vortexing and pipetting were used to disperse the cells prior to serial dilution in PBS. For each sample, 100 μL of undiluted PBS was plated in a lawn and 10 μL of each serially diluted sample is plated on PIA for colony counts. Lactate dehydrogenase (LDH) release from epithelial cells was assayed using the CytoTox 96 nonradioactive cytotoxicity assay.

### *Drosophila melanogaster* colonization model

Colonization of *Drosophila melanogaster* with *P. aeruginosa* strain FRD1 was performed as previously described (54–56). Briefly, Canton S flies were treated with an antibiotic cocktail (erythromycin, vancomycin, and ampicillin at 50 μg/ml) for 3 days and transferred to antibiotic-free Jazz-Mix fly food for 3 days to remove residual antibiotics. On the day of infection, flies were starved for 3 hours prior to being added to vials (10 flies per vial). To infect flies, 1.5mLs of 16-hour FRD1 cultures grown in SCFM2 were pelleted and re-suspended in 100 µL of 5% sucrose (+/- 10 µg/mL tobramycin) and overlayed on a 21-mm Whatman filter that was placed on 5mL of 5% sucrose agar in a plastic vial (FlyBase), followed by the addition of flies. Following 24 hours of oral infection, flies were anesthetized with CO_2_, briefly sterilized with 70% ethanol, and washed 3 times with sterile PBS. Flies were homogenized in PBS and the homogenate was serially diluted and plated on Pseudomonas Isolation Agar to quantify *P. aeruginosa*.

### RNA Extraction and Sequencing

Bacterial cultures were grown in triplicate to exponential phase in SCFM2. Cultures were pelleted, resuspended in Trizol, and cells were lysed with glass beads. RNA was extracted using the Zymo Research Direct-Zol RNA MiniPrep. RNA quality was assessed at the UAB Heflin Genomics Core prior to being processed by SeqCenter for sequencing. Annotation of the genomes was generated using Prokka. Reads were first mapped using STAR and the read counts per annotated gene were determined using featureCounts. Differential gene expressions were then computed using DESeq2 and genes with p-values less than 0.05 and fold changes with absolute changes greater than 1.5 were filtered. The transcriptomics data were deposited in Gene Expression Omnibus under accession number GSE280261.

### Biochemical Assays

All cultures were grown in SCFM2 for all biochemical assays for 24 hours. Alginate was purified from culture supernatants that were dialyzed against distilled water and quantified using the carbazole method as previously described (57). Pyocyanin was extracted using chloroform as previously described (57).

### Statistical analysis

All experiments were conducted using a minimum of 3 biological and 3 technical replicates. Unless otherwise noted, graphs represent sample means ± standard error of the mean (SEM). Analysis of variance (ANOVA) or t-test statistical tests were performed using GraphPad Prism 10 (San Diego, CA).

## Acknowledgements

This work was supported by grants and funds awarded to J.A.S from the National Institute of General Medical Sciences (R35GM142748), the UAB School of Medicine Pittman Scholars Fund, and start-up funds from the UAB Department of Microbiology.

